# Suramin potently inhibits binding of the mammalian high mobility group protein AT-hook 2 to DNA

**DOI:** 10.1101/838656

**Authors:** Linjia Su, Nadezda Bryan, Sabrina Battista, Juliano Freitas, Alyssa Garabedian, Federica D’Alessio, Miriam Romano, Fabiana Falanga, Alfredo Fusco, Lidia Kos, Jeremy Chambers, Francisco Fernandez-Lima, Prem P. Chapagain, Stefan Vasile, Layton Smith, Fenfei Leng

**Author notes:** To whom correspondence should be addressed: Department of Chemistry & Biochemistry, Florida International University, 11200 SW 8^th^ Street, FL 33199.

## Abstract

The mammalian high mobility group protein AT-hook 2 (HMGA2) is a multi-functional DNA-binding protein which plays important roles in tumorigenesis and adipogenesis. Previous results showed that HMGA2 is a potential therapeutic target of anticancer and anti-obesity drugs by inhibiting its DNA-binding activities. Here we report the development of a miniaturized, automated AlphaScreen high throughput screening (HTS) assay to identify inhibitors targeting HMGA2-DNA interactions. After screening the LOPAC1280 compound library, we discovered that suramin, a negatively charged antiparasitic drug potently inhibits the HMGA2-DNA interaction. Our results also show that the inhibition is through suramin binding to the AT-hooks of HMGA2, therefore blocking its DNA binding capacity. Furthermore, we demonstrate that suramin can induce brain tumor stem cells differentiation into cells with neurite-like structures, a process triggered by disrupting HMGA2-DNA interactions. Since suramin has strong antitumor and anti-metastasis activities, our discovery suggests that HMGA2 and HMGA2-like proteins may be the cellular target of this century-old drug.

---

The mammalian high mobility group protein AT-hook 2 (HMGA2) is a multi-functional nuclear protein associated with epithelial-to-mesenchymal transition (EMT) during embryonic development (1). Early studies showed that HMGA2 is related to preadipocyte proliferation and obesity (2–4). For example, *Hmga2* knockout mice were severely deficient in fat cells and developed pygmy phenotype (5). The disruption of *Hmga2* gene dramatically reduced obesity of leptin-deficient mice (*Lep^ob^/Lep^ob^*) (2). These results suggest that HMGA2 is a potential target for the treatment of obesity. HMGA2 is also linked to oncogenesis. Its over and/or aberrant expression leads to the formation of a variety of tumors including benign tumors, such as lipomas (6), uterine leiomyomas (7), and fibroadenomas (8), and malignant tumors, such as lung cancer (9, 10), breast cancer (11, 12), prostate cancer (13), leukemia (14), and melanoma (15–18). Intriguingly, HMGA2 expression level always correlates with the degree of malignancy, metastasis, and a poor prognosis (19, 20), suggesting that this protein is also a therapeutic target of anti-cancer and anti-metastasis drugs (21, 22). Furthermore, HMGA2 is associated with neural and hematopoietic stem cell youth (23, 24), human height (25), and human intelligence (26).

HMGA2 is a small DNA-binding protein consisting of three “AT-hook” DNA binding motifs and a highly acidic C-terminal motif (27). These three “AT-hooks” contain a unique palindromic sequence, PGRGP, each side surrounded by one or two positively charged amino acids, i.e., Lysine or Arginine (28). HMGA2 is an intrinsically disordered protein (IDP (29)). When it binds to AT-rich DNA sequences, the “AT-hook” DNA binding motifs adopt defined structures (30). This disordered-to-ordered conformational transition allows HMGA2 to adapt to different AT-rich DNA sequences and to participate in different nuclear activities, such as transcription (31), DNA replication (32), and DNA repair (33). These results suggest that HMGA2-DNA interactions could be chemically intervened for therapeutic purposes. Utilizing a systematic evolution of ligands by exponential enrichment (SELEX) method, we previously identified two consensus DNA sequences for HMGA2 binding: 5′-ATATTCGCGAWWATT-3′ and 5′-ATATTGCGCAWWATT-3′, where W is A or T (27). This result and a following study (34) suggests that HMGA2 binds and bends specific DNA sequences.

With the identification of HMGA2 as a potential target for the treatment of obesity and cancer, the next step is to search for chemical compounds that prevent HMGA2 binding to its target DNA sequences. For instance, utilizing protein–DNA interaction enzyme-linked immunosorbent assays (PDI-ELISA), we recently found several DNA-binding inhibitors including netropsin that disrupt HMGA2-DNA interactions (35). Intriguingly, our results showed that netropsin strongly inhibited the differentiation of the mouse pre-adipocyte 3T3-L1 cells into adipocytes most likely through a mechanism by which netropsin inhibits HMGA2-DNA interactions (35). However, DNA-binding compounds are usually too toxic to be used as anti-obesity and anticancer drugs. Novel inhibitors of HMGA2-DNA interactions that are not cytotoxic and do not directly bind DNA are required before such approaches can be considered as therapeutic applications. A high-throughput screening (HTS) strategy is needed to identify novel compounds that disrupt HMGA2-DNA binding. To achieve this, we developed an AlphaScreen-based assay of HMGA2 binding to DNA that is amenable to automated HTS in a high-density plate format. Here we report the establishment of the assay, and the identification of suramin, a century-old antiparasitic drug, as a potent inhibitor of HMGA2-DNA interactions.

## Results

### An automated HTS assay for inhibitors of HMGA2-DNA complexes

We previously used a PDI-ELISA assay to screen a small library containing 29 DNA-binding compounds and successfully identified several small molecules that disrupt HMGA2 binding to the minor groove of AT-rich DNA sequences (35). Although this assay performs well in a 96 well plate format, it is not suitable for automated ultrahigh throughput screening of 100,000s of compounds due to the need for streptavidin-coated assay plates and multiple wash steps. To address these limitations, we established a new assay using AlphaScreen technology. The assay entails binding a biotin-labeled DNA oligomer FL814 and His-tagged HMGA2 to streptavidin-coated donor beads and nickel chelate (Ni-NTA) acceptor beads, respectively (Fig. 1A). A series of DNA binding studies were performed to determine the optimal conditions for the AlphaScreen Primary Assay (Fig. S1A-D). After these experiments, 12.5 nM of FL814 and 62.5 nM of HMGA2 were chosen for the assay. The assay tolerated up to 1% DMSO without any significant change in signal. We have previously identified two commercially available compounds that strongly inhibit HMGA2 binding to FL814, netropsin and WP631 (35). These two inhibitors are readily available for purchase and served as positive controls for HMGA2-DNA interaction inhibition in the assay. Results in Figs. 1B and S1E demonstrate that netropsin and WP631 potently inhibit HMGA2-DNA interactions with an IC_50_ of 22 and 48 nM, respectively.

**Figure 1.**
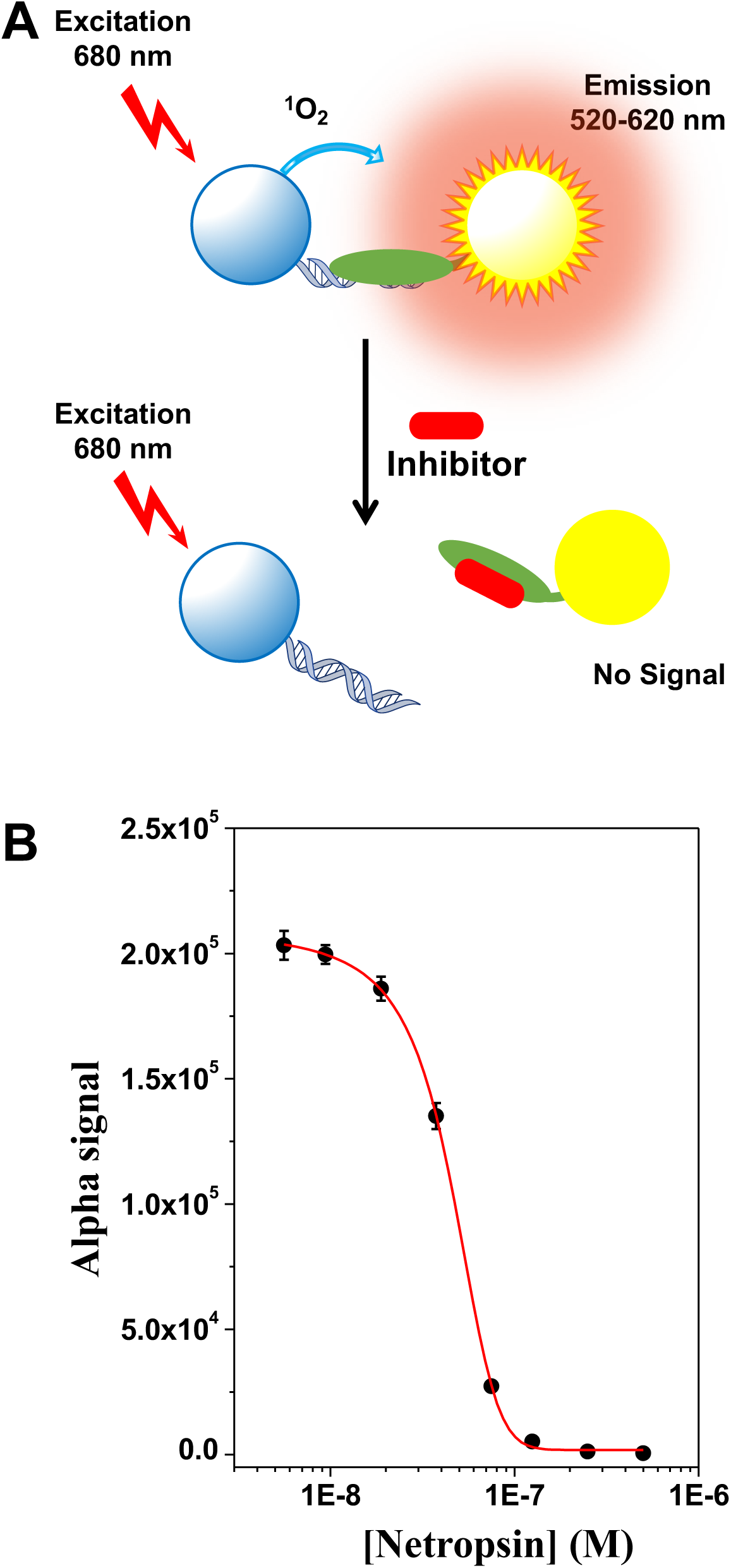
**(A)** The AlphaScreen primary assay for HMGA2-DNA interaction. The biotin-labeled FL814 (double-stranded DNA) and the HIS-tagged HMGA2 (green oval) were immobilized to streptavidin-coated donor beads and nickel chelate (Ni-NTA) acceptor beads, respectively. **(B)** The inhibition of HMGA2 binding to FL814 by netropsin. The curve represents the best fit of a four-parameter logistic that was determined by nonlinear regression. Data points represent mean ±SEM.

In this study, we also developed a LANCE time-resolved fluorescence energy transfer (TR-FRET) assay as a secondary assay for our screening. The His-tag HMGA2 and biotinylated oligomer FL814 were linked to Europium-labeled anti-6×His antibody and APC-labeled streptavidin, respectively. A series of DNA binding studies were performed to determine the optimal conditions for the secondary Assay (Fig. S2). After these experiments, 12.5 nM of FL814 and 62.5 nM of HMGA2 were chosen for the assay, which is the same for the primary assay. The assay tolerated up to 0.25% DMSO without any significant change in signal (Fig. S2C). For a counterscreen assay to exclude those compounds that nonspecifically bind to the protein surface, an AlphaScreen assay of H4 peptide binding to bromodomain-containing protein 4 (BRD4) was adopted (Fig. S2E). Since we are looking for compounds with anti-metastasis or anti-obesity activities, compounds with high levels of cytotoxicity are undesirable. A cytotoxicity assay using the ATPlite Luminescence Assay System and the human epidermoid carcinoma cell line, A431, was developed to eliminate compounds that exhibit cytotoxicity. Compounds that potently and selectively inhibit HMGA2 binding to FL814, and display a LC_50_ >50 µM are will be prioritized for further characterization. Those that do not will be excluded.

### Screen the LOPAC1280 compound library

With the establishment of the miniaturized, automated HTS assays, we screened the Sigma LOPAC1280 collection of pharmacologically active chemical compounds at a final concentration of 5 µM (Fig. S3A). Figs. 2 & S3B and Table S1 show our results and parameters of the AlphaScreen primary HTS assay. The following 16 compounds showed ≥50% inhibition of HMGA2-FL814 interactions: cisplatin, cDPCP, carboplatin, mitoxantrone, Ro 90-7501, aurintricarboxylic acid, GW5047, indirubin-3’-monoxime, 6-hydroxy-DL-DOPA, methyl-3,4-dephostatin, tyrphostin 51, (2’Z,3’E)-6-bromoindirubin-3′-oxime, reactive blue-2, JFD00244, steviol, and suramin. These 16 compounds were cherry picked and then subjected to testing in the LANCE TR-FRET secondary assay. The following seven compounds demonstrated ≥50% inhibition of HMGA2-DNA interactions in both HTS assays: cisplatin, cDPCP, carboplatin, mitoxantrone, Ro 90-7501, aurintricarboxylic acid, and suramin. Dry powders of these seven compounds were purchased for additional testing in these assays. The identity of the compounds was confirmed by LC/MS, and fresh stock solutions (10mM) were prepared in 100% DMSO. We next performed a series of titration experiments and determined the potencies (IC_50_) values of these seven hits in both the primary AlphaScreen assay, and the secondary LANCE TR-FRET assay, as well as the counter screen assay, and cytotoxicity assay. Table S2 summarizes our results. Cisplatin, cDPCP, carboplatin, and mitoxantrone are known DNA-binding agents and likely represent compounds that inhibit HMGA2-DNA interactions by binding to the AT-rich DNA sequence of FL814. Additionally, these DNA-binding compounds may also inhibit other essential cellular functions, which prevent them from further investigation. Intriguingly, although Ro 90-7501, aurintricarboxylic acid, and suramin do not bind to DNA due to their chemical properties, these three compounds strongly inhibit HMGA2-DNA interactions. Of particular interest is suramin, a highly negatively charged antiparasitic drug (36) (Fig. S4) that potently inhibits HMGA2-FL814 integrations with an IC_50_ of 2.58 µM (Fig. 3). Additionally, suramin did not inhibit H4 peptide binding to BRD4 in the counter screen assay (Fig. 3C) and is not cytotoxic to human A-431 cells (Fig. 3D and Table S2). As a final validation of suramin as a “hit” from the HTS, we confirmed its inhibitory effect on HMGA2 interactions with DNA using the PDI-ELISA assay (**Table 1**). Thus, suramin meets all criteria that we set for the identification of novel; HMGA2-DNA inhibitors.

**Figure 2.**
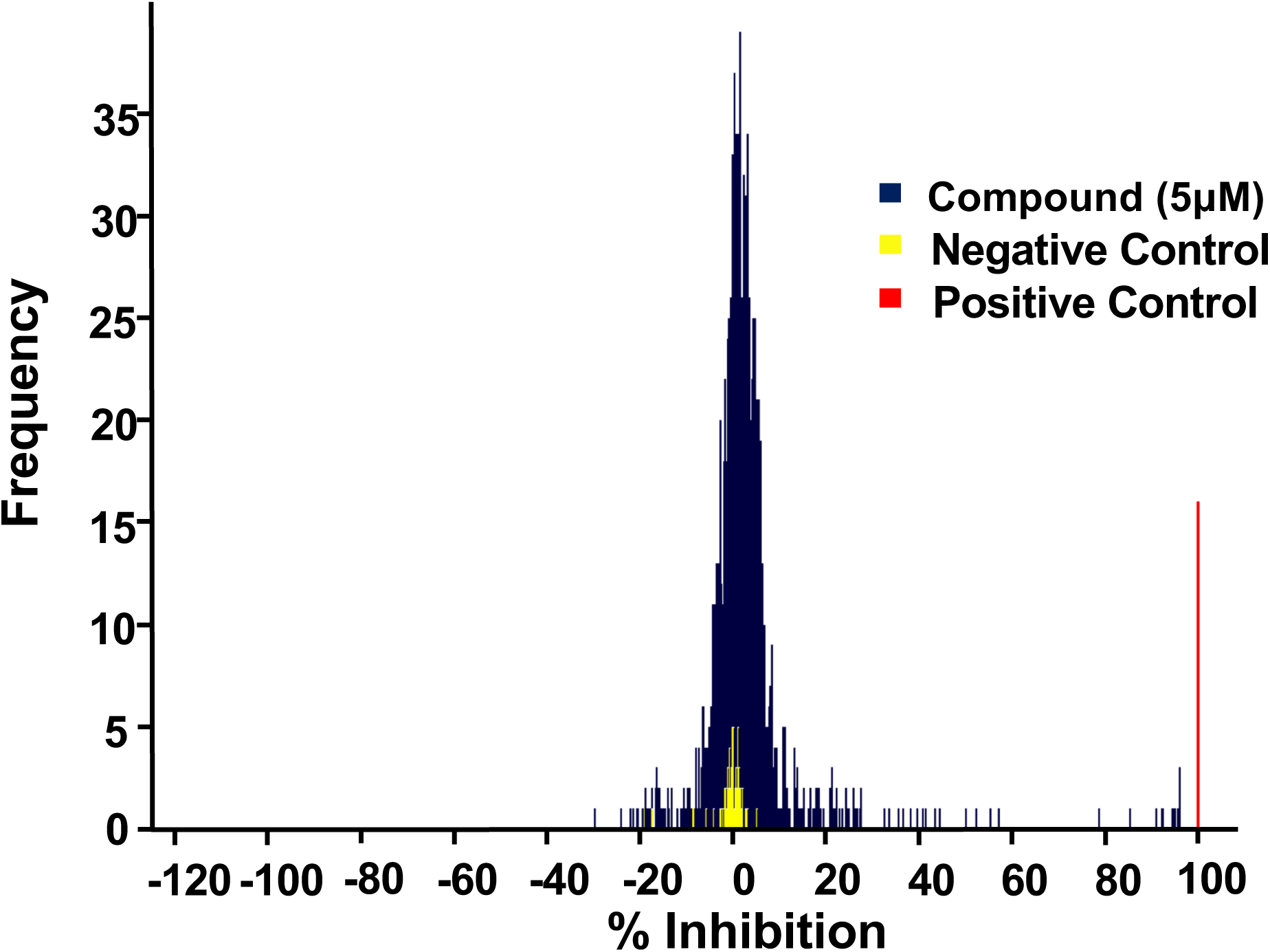
HMGA2-DNA pilot screens using the Sigma LOPAC1280 compound library. Netropsin was used as positive controls.

**Figure 3.**
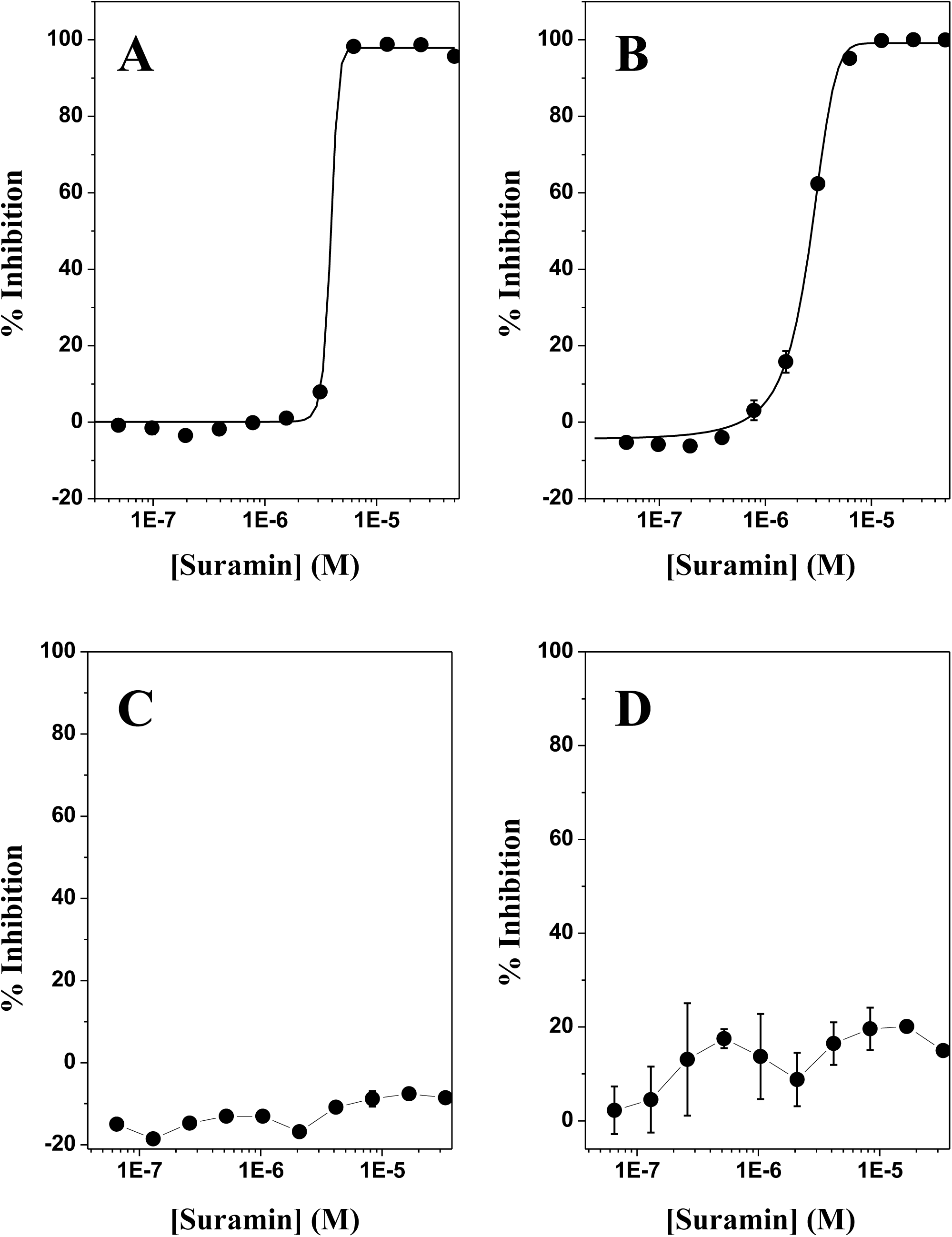
The discovery of suramin as a specific and potent inhibitor of HMGA-DNA interactions. **(A)** The AlphaScreen Primay assay with IC_50_ of 2.6 μM. **(B)** The TR-FRET LANCE secondary screen. **(C)** The couterscreen assay using the BRD4 AlphaScreen Assay. **(D)** The cytotoxicity assay. All assays were described in Materials and Methods. The standard deviation was calculated according to three independent experiments. The curve represents the best fit of a four parameter logistic that was determined by nonlinear regression. Data points represent mean ±SEM.

**Table 1.** Tightly binding of suramin and analogues to HMGA2 and ATHP3.

| Suramin and analogous | IC50 ($\mu\text{M}$) | HMGA2 | | ATHP3 | |
| --- | --- | --- | --- | --- | --- |
| | | $K_a(\text{M}^{-1})$ | Binding site | $K_a(\text{M}^{-1})$ | Binding site |
| Suramin | $2.78 \pm 0.10$ | $4.08 \pm 0.92 \times 10^6$ | 2.81 | $4.06 \pm 0.32 \times 10^7$ | 0.98 |
| | | $4.34 \pm 2.16 \times 10^4$ | 2 | | |
| NF449 | $0.43 \pm 0.18$ | $5.53 \pm 6.14 \times 10^7$ | 0.89 | $1.10 \pm 7.35 \times 10^{11}$ | 0.86 |
| | | $8.17 \pm 1.33 \times 10^5$ | 1.63 | | |
| NF110 | $0.87 \pm 0.039$ | $2.04 \pm 0.48 \times 10^8$ | 2.06 | $2.17 \pm 0.17 \times 10^7$ | 1.01 |
| | | $7.16 \pm 0.67 \times 10^5$ | 1.77 | | |
| NF546 | $5.49 \pm 2.81$ | $1.48 \pm 0.47 \times 10^7$ | 1.83 | $4.28 \pm 0.47 \times 10^6$ | 0.81 |
| | | $3.37 \pm 8.81 \times 10^5$ | 1.98 | | |
| NF340 | $6.95 \pm 9.36$ | $7.17 \pm 4.68 \times 10^6$ | 2.12 | $1.89 \pm 0.16 \times 10^7$ | 0.8 |
| | | $2.27 \pm 0.57 \times 10^5$ | 0.3 | $1.77 \pm 0.06 \times 10^5$ | 1.06 |
| NF023 | $10.63 \pm 0.46$ | $2.47 \pm 4.53 \times 10^6$ | 1.74 | $9.75 \pm 3.47 \times 10^6$ | 1 |

### Suramin and analogues strongly bind to HMGA2 and ATHP3

We next sought to determine the mechanism by which suarmin inhibits HMGA2-DNA interactions. Since suramin carries 6 negative charges (Fig. S4), it should not bind to DNA due to the fact that DNA is highly negatively charged. Instead, it should bind to HMGA2 because HMGA2 is positively charged (37). ITC studies revealed that suramin physically interacts with HMGA2 (Fig. 4A and Table 1). These studies revealed that there are two types and a total of five suramin binding sites on HMGA2. The first type of three suramin binding sites has a binding constant of 4.08 ± 0.92 × 10^6^ M^-1^ with the following thermodynamic parameters: ΔG, -9.02 kcal/mol; ΔH -14.58 kcal/mol; and –TΔS, 5.56 kcal/mol. The second type of two suramin binding sites has a binding constant of 4.34 ± 2.16 × 10^4^ M^-1^ with the following thermodynamic parameters: ΔG, -6.33 kcal/mol; ΔH -6.57 kcal/mol; and –TΔS, 0.24 kcal/mol. These five suramin binding sites of HMGA2 were confirmed by our mass spectrometric experiments at high suramin to HMGA2 ratios (Fig. S5). Because HMGA2 contains three highly positively charged “AT-hook” DNA binding motifs, it is reasonable to assume that suramin strongly binds to these highly positively charged motifs through charge-charge interactions. Our ITC experiments of suramin titrating into an “AT-hook” peptide 3 (ATHP3) solution confirmed this hypothesis: suramin binding to ATHPs has a binding constant of 4.06±0.32×10^7^ M^-1^ and 1:1 binding molar ratio (Fig. 4B and Table 1). The following are its binding thermodynamic parameters: ΔG, - 10.38 kcal/mol; ΔH -17.13 kcal/mol; and –TΔS, 6.75 kcal/mol.

**Figure 4.**
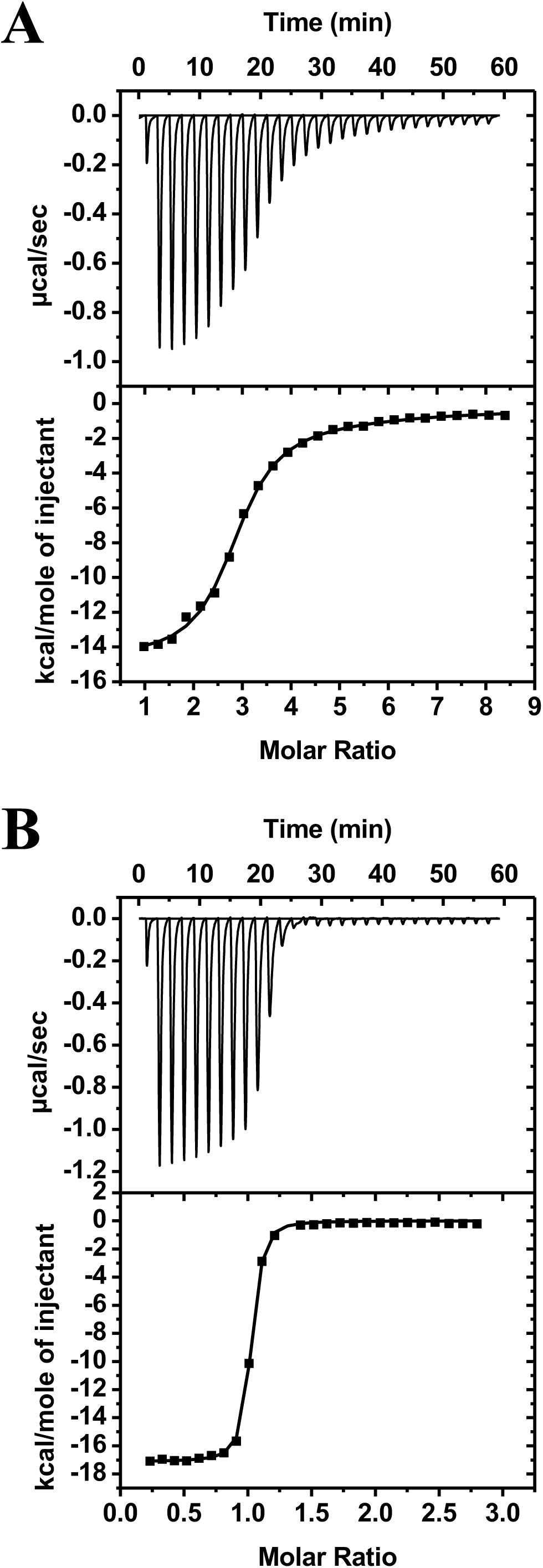
Sample raw data from isothermal titration calorimetry (ITC) experiments for the titration of suramin to HMGA2 **(A)** and ATHP3 **(B)**. ITC experiments were performed according to conditions as described in Materials and Methods. The ITC data were fit using the software supplied by the manufacturer to yield thermodynamic parameters.

Next, we examined the structure activity relationship (SAR) of suramin using several suramin analogues (Figs. S4, S6, and S7). As shown in Table 1, all of these analogues except sodium 1-naphthalenesulfonate potently inhibit HMA2-DNA interactions with IC50 values ranging from 0.43 to 10.63 µM. Likewise, all these chemical compounds except sodium 1-naphthalenesulfonate strongly bind to both HMGA2 and ATHP3. Intriguingly, although the negative charge is important, it is not the only parameter that determines their inhibition potency. For example, NF110 only carries 4 negative charges. However, it strongly inhibits HMGA2-DNA interactions with an IC50 of 0.87 ± 0.04 µM. In contrast, although NF023 has 6 negative charges, its inhibition IC50 was determined to be 10.63 ± 0.46 µM, 10-fold higher than that of NF110. These results suggest that both the charge and structure are very important for the inhibition of HMGA2-DNA interactions by these related compounds.

### Suramin induces the differentiation of brain tumor stem cells and inhibits aggressiveness of B16F10 melanoma cells

As mentioned above, over and/or aberrant expression of HMGA2 is associated with the formation of a variety of benign and malignant tumors (21). For instance, previous data have shown that HMGA proteins including HMGA2 are highly expressed in glioblastomas and glioblastoma-derived brain tumor stem cells (BTSCs) (38), where they play pivotal roles in regulating self-renewal, differentiation and symmetric/asymmetric division (38, 39). Therefore, BTSCs are an optimal cell assay system to evaluate the biological effect of suramin mediated by HMGA2. Here, we exposed two BTSC lines (BTSC#83 and BTSC#30p) with different concentrations of suramin. Consistent with the cytotoxicity studies (Table S2), up to 200 μM of suramin did not significantly affect the cell growth of these two cell lines (Fig. S8A). Intriguingly, in the presence of 100 µM and 200 µM (Fig. 5A) of suramin, cells were stimulated to adhere to the bottom of the well and extend neurite-like structures (as in BTSC#83; Fig. 5A, top panel) or acquire flat epithelial morphology (as in BTSC#30p; Fig. 5A, bottom panel), suggesting induction of differentiation in both BTSC cell lines. Western blotting and qRT-PCR experiments showed that suramin slightly reduced the expression of HMGA2 and HMGA1in both cell lines (Fig. 5B and C). Nevertheless, it significantly reduced the expressions of ID2, a cell-stemness marker and SNAIL & TWIST, two EMT markers (Fig. 5C). In contrast, suramin increased the expression of NUMB, a cell-fate determinant usually upregulated in differentiating cells. Interestingly, our results showed that the downregulation of ID2, SNAIL, and TWIST and upregulation of NUMB is proportional to the expression level of HMGA2 for these two BTSCs. In other words, suramin has a greater effect on these protein markers (ID2, SNAIL, TWIST, and NUMB) in BTSC#30p, which has a higher HMGA2 expression level, than that in BTSC#83 (Fig. 5C). These results suggest that suramin targets HMGA2-DNA interactions in these two BTSC cell lines to induce their differentiation.

**Figure 5.**
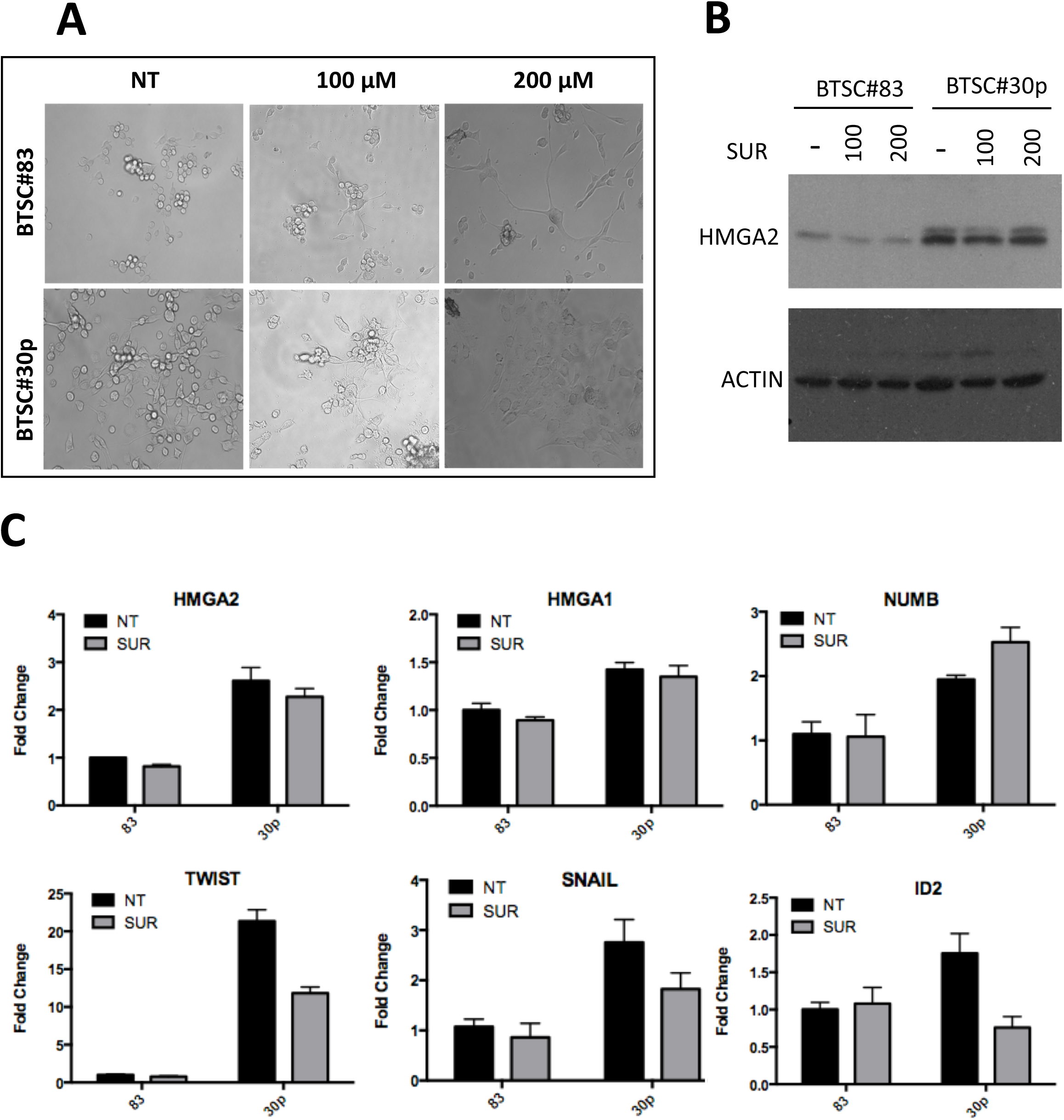
Suramin induces the differentiation of brain tumor stem cells (BTSCs) 83 and 30p. (A) Light Microscopy of BTSC#83 and BTSC#30p treated with various concentrations of suramin for 6 days, compared to non-treated (NT) cells. (B) Western blot analysis of HMGA2 expression after the 6-day treatment of suramin. (C) Expression of HMGA2 and EMT markers in suramin-treated BTSCs. qRT-PCR analyses in BTSC#83 and BTSC#30p treated with suramin 100 µM for 48 hrs, compared to non-treated cells. Fold changes are normalized for actin expression. Means and standard deviations are calculated from three independent experiments.

Melanoma is the least common but the most aggressive type of skin cancer (40). Previous studies showed that HMGA2 is highly expressed in primary & metastatic melanomas (41) and also in various melanoma cell lines (15). Since overexpression is associated with melanoma progression and prognosis, it may serve as an independent predictor for melanoma survival (16). Additionally, HMGA2 is related to the EMT and migratory potential of melanoma cells (42). Therefore, we next examined how suramin affects migratory behavior and the expression of E-cadherin of B16F10 melanoma cell line, a highly aggressive murine tumor cell line used as a model for the study of metastasis and solid tumor formation (15). Increased migratory behavior is one of the prototypic phenotypes associated with aggressiveness (43). We assessed the migratory behavior of B16F10 using a wound-healing assay. Our results demonstrated that 100 µM Suramin is sufficient to significantly inhibit the migratory behavior of B16F10 melanoma cells (Fig. S8B). This result is consistent with previously published results (44). Loss of E-cadherin is also associated with the aggressive phenotype of cancer cells (45). Interestingly, E-cadherin is frequently down-regulated in melanoma (46). The treatment of B16F10 melanoma cells with 100 µM Suramin significantly increased the expression level of E-cadherin when compared to non-treated cells (Fig. S8C). These results suggest that suramin is a potent agent to block the aggressive features of the mouse melanoma cell line. Consistent with our BTSCs’ results, suramin does not significantly affect the expression of HMGA2 and however greatly decreases the expression of SNAIL and SLUG assessed by qRT-PCR (data not shown).

## Discussion

In this article, we report the development of a miniaturized, automated AlphaScreen HTS assay to identify small molecule inhibitors targeting HMGA2-DNA interactions. This HTS assay has excellent screening parameters with a Z’ and S/B ratio at 0.83 and 438, respectively (Table S1). After screening the LOPAC1280 compound library, we found that suramin, a highly negatively charge anti-parasitic drug that is used to treat African sleeping sickness and river blindness (47, 48), is a potent inhibitor of HMGA2-DNA interactions. Further, our results show that suramin’s inhibition of HMGA2-DNA interactions stems from its binding to HMGA2 with high affinity, particularly to ATHPs. Our molecular modeling studies confirmed this hypothesis (Fig. 6). The suramin-ATHP3 complex shows that the suramin sulfonated groups are close to the Arg and Lys residues, providing ionic/hydrogen-bond interactions. The ring structures at the two ends of suramin are folded to embrace the central proline ring from either side of the residue.

**Figure 6.**
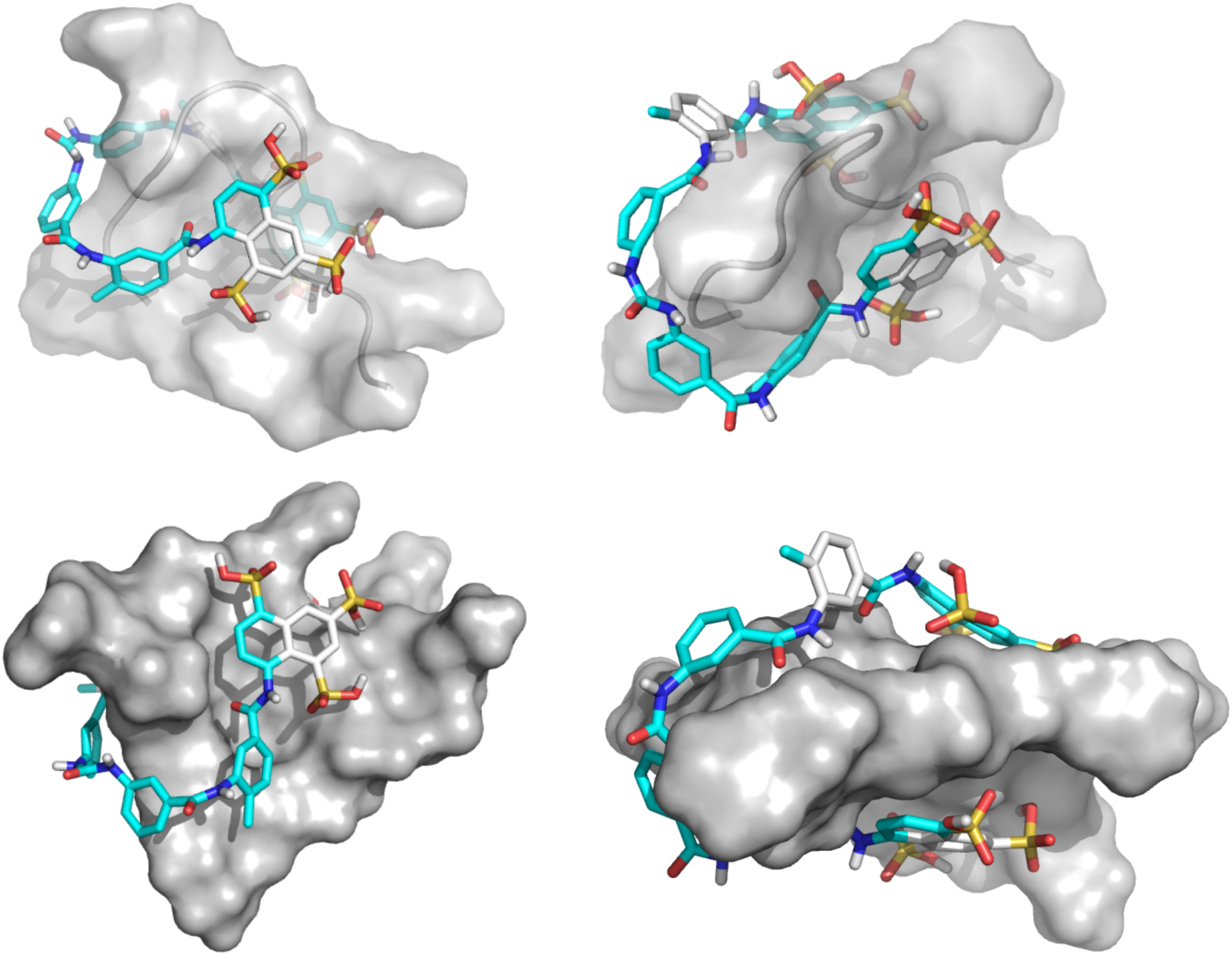
Structure of the suramin-ATHP3 complex obtained by molecular docking and simulations.

DNA-binding proteins, such as transcriptional factors are excellent targets for anticancer therapy (49). Indeed, several clinically important anticancer drugs, such as doxorubicin and cisplatin (50, 51), bind to DNA, disrupt protein-DNA interactions, and as a result prevent DNA-binding proteins including transcriptional factors from binding to their target DNA sites (52). However, a major disadvantage of these anticancer drugs is that they bind to DNA nonspecifically. Consequently, they kill cancer cells as well as normal cells causing serious side effects. Therapeutically active compounds that specifically block oncogenic transcriptional factors from binding to their DNA recognition sites would be superior medicines. Such medicines would be expected to have fewer, less severe side effects, thereby potentially increasing the dose and duration of chemotherapy and perhaps improving treatment outcomes as well as quality of life. Transcription factors are widely considered to be “undruggable” because they usually do not have enzymatic activities, and lack deep pockets to accommodate drug-like molecules (53). Further, the lack of HTS assays to identify inhibitors from small molecule repositories also contributes to the perceived undruggable nature of transcriptional factors (35). Our results refute these notions, at least with respect to the potential drugability of HMGA2. Previously, we developed protein–DNA or protein–RNA interaction enzyme-linked immunosorbent assays (PDI-ELISA or PRI-ELISA) to identify inhibitors targeting specific protein-nucleic acids interactions (35). These methods are versatile and can be applied to any nucleic acid binding proteins as long as an antibody is available. In fact, one may use tagged proteins, such as His-tagged proteins so that antibodies against His-tag can be used in PDI-ELISA or PRI-ELISA. Regardless, as we pointed out above, because streptavidin-coated 1536-well plates are expensive, PDI-ELISA or PRI-ELISA cannot be configured into a miniaturized, automated ultrahigh throughput screens in a 1536 well plate format. Additionally, too many washing steps were used, which make PDI-ELISA or PRI-ELISA a “lengthy” procedure and unsuitable for automated HTS studies. Here we showed that AlphaScreen assays could be utilized to screen for inhibitors against HMGA-DNA interactions by using His-tagged HMGA2 linked to nickel chelate (Ni-NTA) acceptor beads. The use of AlphaScreen HTS assays has several advantages. Because His-tag can be added to most of DNA-binding proteins and usually does not interfere with their DNA binding activities, AlphaScreen assays can be used to screen and identify inhibitors targeting most DNA-binding proteins. Another advantage is that AlphaScreen HTS assays are cost-effective. For instance, in this study, only one 1536-well plate was used in the screening of the LOPAC1280 compound library. It is anticipated that AlphaScreen HTS assays using His-tagged DNA binding proteins will be utilized to identify inhibitors for other DNA-binding proteins in the future.

The most intriguing discovery of this HTS study is that suramin potently inhibits HMGA2-DNA interactions likely through a mechanism by which suramin tightly binds to highly positively charged ATHPs. This is not surprising because suramin, a polysulphonated naphylurea, carries 6 negative charges (Fig. S4). As shown in Fig. 6, charge-charge interactions and hydrogen bonding between the suramin sulfonated groups and Arg/Lys residues play critical roles in the binding of suramin to ATHPs. This inhibition mechanism is different from that of netropsin and other DNA minor groove binders that prevent HMGA2 from binding to the minor groove of AT-rich DNA sequences (22, 35). In fact, suramin is the first chemical compound that was found to tightly bind to the intrinsically disordered protein (IDP) HMGA2 and the “unstructured” DNA binding motif ATHPs. This discovery leads to an important and interesting question: can suramin bind to HMGA2 or HMGA2-like proteins *in vivo*? Although suramin is a century-old drug and was synthesized in the 1920s by Bayer to treat human African trypanosomiasis (HAT) (54), the mechanism or mode of action is still unknown (55). Suramin is usually administered by intravenous injection due to the poor intestinal absorption (36) and binds to serum proteins, such as albumin and low-density lipoprotein (LDL), immediately after administration (36). It is believed that the parasite takes up the drug through receptor-mediated endocytosis of the protein-bound drug (56). Suramin is highly effective against blood-stream forms of the parasite but not very active against procyclic trypanosomes (57). Since the bloodstream forms of Trypanosoma brucei lacks a functional mitochondrion and entirely depend on glycolysis for their energy needs, this led to the hypothesis that the glycolytic pathway of the parasite is the target of the drug (58). However, so far there is no direct evidence to support this hypothesis (36). Another interesting feature of Trypanosoma brucei is that its mitochondrion contains the so-called kinetoplast DNA (kDNA), comprising of ∼73% AT base pairs (59). Recent studies showed that certain DNA minor groove binding compounds were able to enter the mitochondrion of Trypanosoma brucei, bind to AT sequences of kDNA and, as a result, cause cell death (60). It was suggested that these minor groove binding compounds may disrupt the functions of kDNA binding proteins, such as the TbKAP6 protein that is essential for kDNA replication and maintenance and also for cell viability (61). TbKAP6 contains two HMG boxes and binds to DNA minor groove (61). Interestingly, this protein carries several highly positively charged motifs similar to ATHP (61). Together with the findings reported here, we hypothesize that suramin could tightly bind to these positively charged motifs in TbKAP6 and prevent its binding to kDNA. Further studies are needed to support this hypothetic mechanism.

Other important functions of suramin are its anticancer and anti-metastasis activities (62, 63). Although suramin’s anticancer and anti-metastasis activities have been studied extensively (62, 63), the mechanism is still unknown. Possible mechanisms include inhibiting important enzymes, such as cullin-RING E3 ubiquitin ligases (64), protein-tyrosine phosphatases (65), and blocking various growth factors binding to their receptors (62). The discovery of suramin tightly binding to HMGA2 and potently inhibiting HMGA2-DNA interactions provides a new insight into its anti-cancer and anti-metastasis functions since the expression level of HMGA proteins including HMGA1 and HMGA2 is associated with metastasis and poor prognosis for many cancer types (21). Possibly, binding of suramin to HMGA2 prevents the protein from binding to the target DNA sequences and therefore inhibits its functions in tumorigenesis. Suramin-induced differentiation of BTSCs is consistent with this hypothesis (Fig. 5A). Suramin does not significantly affect HMGA2 expression in both BTSCs (Fig. 5B and C) and however decreases the expression of SNAIL, TWIST, and ID2, transcriptional factors associated with tumorigenesis and metastasis (66). Importantly, the decrease of these three transcriptional factors is correlated with HMGA2 expression level, suggesting that suramin binds to HMGA2 and inhibits HMGA2-DNA interactions inside these BTSCs. Additional evidence to support this hypothesis that suramin inhibits HMGA2-DNA interactions in cancer cells stems from our study using B16F10 melanoma cells. Previously it was shown that suramin alone or combined with pentoxifilline inhibited B16F10 melanoma metastasis in mouse models (44). At low dose concentrations, suramin inhibits the migratory behavior and stimulates the E-cadherin expression of B16F10 melanoma cells (Fig. S8). Intriguingly, microRNA let7a inhibits migration of B16F10 melanoma cells most likely through a mechanism by which it down-regulates HMGA2 expression (17), suggesting that the anti-metastasis activities of suramin result from its tightly binding to HMGA2 or HMGA2-like oncoproteins. Because HMGA2 is associated with tumorigenesis and metastasis for a variety of tumors, future studies of surmain’s anti-tumor and anti-metastasis activities should focus on those tumors that over and/or abnormally express HMGA2. For instance, a recent study using RNA sequencing and survival data of 430 acute myeloid leukemia (AML) patients showed that the high-level expression of HMGA2 is a predictor of poor clinical outcomes for both cancer relapse and patient survival after the standard AML treatment including the use of etoposide or doxorubicin, topoisomerase II poisons (14). *Ex vivo* studies (67) suggested that the poor clinical outcomes of AML with high HMGA2 expression result from the protective role of HMGA2 in etoposide or doxorubicin-induced double-stranded DNA breaks through topoisomerase II poisoning. A combination therapy of etoposide or doxorubicin with suramin may be more effective to treat these AML patients with high expression of HMGA2.

Finally, our results may add additional insights into a small randomized clinical trial showing that suramin is effective to treat autism spectrum disorder (ASD) patients (68). This clinical trial was based on a hypothesis that purinergic signaling plays a critical role in the ASD pathogenesis and suramin blocks this signaling pathway by inhibiting purinergic receptors (69). We showed here that suramin promotes the production of NUMB (Fig. 5C), a transcriptional factor involved in neuronal differentiation by inhibiting HMGA2-DNA interactions and thereby induces the differentiation of BTSCs into neuron-like cells (Fig. 5A). This result suggests that a different, as yet unidentified, mechanism may also contribute to the effectiveness of suramin as a potential drug to treat ASD.

## Materials and Methods

### Materials

The His-tagged mammalian HMGA2 was purified using a Ni-NTA agarose column followed by an SP Sepharose fast flow column as described previously (37). An extinction coefficient of 5,810 M^-1^ cm^-1^ at 280 nm was used to determine its concentration (37). Biotin-labeled DNA oligomer FL814 carrying a specific SELEX binding site of HMGA2 (27, 35) was purchased from Eurofins MWG Operon, Inc. AlphaScreen histidine (nickel chelate) detection kits containing nickel chelate acceptor beads and streptavidin donor beads (#6760619M), LANCE Ultra ULight-anti-6xHIS (#TRF0105) and LANCE Eu-W1024 Streptavidin (#AD0062), as well as 1536 well Optiplates plates (#6004290) were purchased from Perkin Elmer. Suramin and NF023 were purchased from MilliporeSigma, Inc. Sodium 1-Naphthalenesulfonate was obtained from TCI America, Inc. NF110, NF340, NF449, and NF546 were purchased from Tocris Biosciences. His-tagged BRD4-BD1 was purchased from BPS Bioscience (#31042). Pre-acetylated Biotin-Histone 4 Peptide was purchased from AnaSpec (#64989-025). Bovine serum albumin fraction V (#A7888), CHAPS (#C3023-25G), and LOPAC1280 library (#LO1280) were purchased from Sigma Aldrich. Human epidermoid carcinoma A-431 cell line was purchased from the ATCC (ATCC-CRL-1555). DMEM media was purchased from Thermo Fisher (Gibco #11995), 100×Penicillin/Streptomycin solution (#30-002-CI), DPBS (#21-031-CV), 0.25% Trypsin in HBSS (#25-050-CI), 200 mM L-glutamine (#25-005-CI), white high base 1536 well plates (#4570), and sterile white high base 1536 well plates (#4571) were purchased from Corning. The control compound MG-132 was purchased from Promega (#G932B). ATP-Lite 1-step was purchased from Perkin Elmer (#6016739).

### HMGA2 AlphaScreen HTS assay

Using a Labcyte Echo 555 acoustic dispenser 5 nL of DMSO were added to columns 1-4 and 45-48 of a white 1536 well plate, while 5 nL of 2 mM compounds in DMSO were added to columns 5-44. Using a Beckman BioRAPTR FRD bulk reagent dispenser 1 µL of assay buffer (30 mM Citrate, 300 mM NaCl, and 0.005% Tween 20) was added to columns 1 and 2. Next 1 µL of assay buffer containing 125 nM HMGA2 was added to columns 3-48 with a Beckman BioRAPTR FRD bulk reagent dispenser. Finally, 1 µL of assay buffer containing 25 nM FL814 was added to every well of the plate using a Beckman BioRAPTR FRD bulk reagent dispenser. The plate was then centrifuged at 200×g for 1 minute. After 30 minutes at room temperature, 2 µL of bead buffer (10 mM HEPES, 300 mM NaCl and 0.005% Tween 20) containing 20 µg/mL anti-6xHIS acceptor beads and 20 µg/mL streptavidin donor beads were dispensed into every well using a Beckman BioRAPTR FRD bulk reagent dispenser. The plates were then centrifuged at 200×g for 1 minute. After 1 hour at room temperature the plates were read on a Perkin Elmer Envision multimode plate reader in AlphaScreen mode.

### HMGA2 Lance assay

Using a Labcyte Echo 555 acoustic dispenser 5 nL of DMSO were added to columns 1-4 and 45-48 of a white 1536 well plate, while 5 nL of 2 mM compounds in DMSO were added to columns 5-44. Using a Beckman BioRAPTR FRD bulk reagent dispenser 1 µL of assay buffer (10 mM Tris, 300 mM NaCl and 0.005% Tween 20) was added to columns 1 and 2. Next, 1 µL of assay buffer containing 125 nM HMGA2 was added to columns 3-48 with a Beckman BioRAPTR FRD bulk reagent dispenser. Finally, 1 µL of assay buffer containing 25 nM FL814 was added to every well of the plate using a Beckman BioRAPTR FRD bulk reagent dispenser. The plate was then centrifuged at 200×g for 1 minute. After 30 minutes at room temperature 1 µL of assay buffer containing 250 nM LANCE Ultra ULight-anti-6xHIS and 1 µL of assay buffer containing 12 nM LANCE Eu-W1024 Streptavidin was dispensed into every well using a Beckman BioRAPTR FRD bulk reagent dispenser. The plates were then centrifuged at 200×g for 1 minute. After 1 hour at room temperature the plates were read on a Perkin Elmer Envision multimode plate reader in TR-FRET mode (excitation @340 nm, first emission at @665nm, second emission at 615nm).

### BRD4 AlphaScreen Assay

Using a Labcyte Echo 555 acoustic dispenser 5 nL of DMSO were added to columns 1-4 and 45-48 of a white 1536 well plate, while 5 nL of 2mM compounds in DMSO were added to columns 5-44. Using a Beckman BioRAPTR FRD bulk reagent dispenser 1 µL of assay buffer (50 mM HEPES, 100 mM NaCl, 0.1% BSA and 0.0005% CHAPS) was added to columns 1 and 2. Next, 1 µL of assay buffer containing 50 nM BRD4 was added to columns 3-48 with a Beckman BioRAPTR FRD bulk reagent dispenser. Finally, µL of assay buffer containing 50 nM peptide was added to every well of the plate using a Beckman BioRAPTR FRD bulk reagent dispenser. The plate was then centrifuged at 200×g for 1 minute. After 60 minutes at room temperature 2 µL of assay buffer containing 20 µg/mL anti-6xHis acceptor beads and 20 µg/mL streptavidin donor beads was dispensed into every well using a Beckman BioRAPTR FRD bulk reagent dispenser. The plates were then centrifuged at 200×g for 1 minute. After an overnight incubation at room temperature the plates were read on a Perkin Elmer Envision multimode plate reader in AlphaScreen mode.

### Cell viability assay

A-431 cells were grown in media (DMEM + 10% FBS + 1×Penicillin/Streptomycin + 2 mM L-glutamine) at 37°C in a humidified incubator with 5% CO_2_ to 70% confluency. They were then washed with DPBS and trypsinized. For the assay the cells were resuspended in media (DMEM + 10% FBS + 1×Penicillin/Streptomycin + 2 mM L-glutamine) at 125,000 cells/mL. Using a Thermo multidrop combi bulk reagent dispenser 4 µL of cell suspension (500 cells) were dispensed into every well of a 1536 well plate. The plates were centrifuged at 200×g for 1 minute, relidded and incubated overnight at 37°C with 5% CO_2._ The next day using a Labcyte Echo 555 acoustic dispenser 5 nL of MG-132, final in well concentration of 25 µM, were added to columns 1-2, 5 nL of DMSO were added to columns 3-4 and 45-48 and 5 nL of 2 mM compounds in DMSO were added to columns 5-44. The plates were then centrifuged at 200×g for 1 minute, relidded and returned to the incubator. After 48 hours 4 µL of room temperature ATP-Lite were added with a Beckman BioRAPTR FRD bulk reagent dispenser. The plates were centrifuged at 200×g for 1 minute and then incubated at room temperature. After 10 minutes the plates were read using a Perkin Elmer Viewlux multimode plate reader in luminescence mode.

### Protein-DNA interaction ELISA (PDI-ELISA) assay

The PDI-ELSA assays were described previously (35) and used to determine the inhibition IC_50_ against HMGA2-DNA interactions. The apparent inhibitory IC_50_ values were obtained using the following equation: 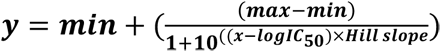, where x, y, max, and min represent the inhibitor’s concentration, the inhibition level, the maximum inhibition value, and the minimum inhibition value, respectively.

### Isothermal titration calorimetry (ITC)

ITC experiments were conducted using a VP-ITC titration calorimeter (MicroCal, Inc., Northampton, MA) interfaced to a PC. Origin 7.0, supplied by the manufacturer was used for data acquisition. For a typical ITC experiment, the titration was set up so that 10 µL of 0.2 mM suramin or analogs was injected every 120 seconds, up to a total of 29 injections, into an HMGA2 sample (1.44 mL of 5 µM) or ATHP3 sample (1.44 mL of 15 µM) in the sample cell in 10 mM Tris-HCl pH 8.0 and 1 mM EDTA. The heat liberated or absorbed is observed as a peak corresponding to the power required keeping the sample and reference cells at identical temperatures. The peak produced during the injection is converted to heat output by integration and corrected for cell volume and sample concentration. Control experiments were also carried out to determine the contribution of the heats of dilution arising from (1) suramin or analogs into buffer and (2) buffer into HMGA2 or ATHP3. The net enthalpy for the titration reaction was determined by subtraction of the component heats of dilution.

### Native Mass Spectrometry

A custom-built nano electrospray unit (nESI) was coupled to a Maxis Impact HD Q-TOF mass spectrometer (Bruker, Billerica, MA) for all the native mass spectrometry analysis. Quartz glass capillaries (O.D.: 1.0 mm and I.D.: 0.70 mm) were pulled utilizing a P-2000 micropipette laser puller (Sutter Instruments, Novato, CA) and loaded with 10 µL aliquot of the sample solution. Sample solutions consisted of 1-10 µM HMGA2 in 10 mM ammonium acetate solution at physiological pH (pH = 6.7). For the observation of the HMGA2-Ligand complexes, a 1:1, 1:3 and 1:10 ratio of 5 µM concentration of the HMGA2 and Ligand (suramin) were prepared in 10mM ammonium acetate and let it rest for 10 minutes prior infusion. A typical nESI source voltage of +/-600-1200 V was applied between the pulled capillary tips and the MS instrument inlet. Ions were introduced via a stainless-steel tube (1/16 x 0.020’’, IDEX Health Science, Oak Harbor, WA) held at room temperature into the TIMS cell. Solvents, methanol, and ammonium acetate salts utilized in this study were analytical grade or better and purchased from Fisher Scientific (Pittsburgh, PA). A Tuning Mix calibration standard (G24221A) was obtained from Agilent Technologies (Santa Clara, CA) and used as received. Mass spectra were processed using Bruker Compass Data Analysis version 5.1 (Bruker Daltonik GmbH).

### Cell cultures and growth curves

BTSC#83 and BTSC#30p and their culture conditions have been described (38). For growth curves, suspension cultures were mechanically disaggregated and filtered through 37 µm filters (StemCell Technologies, Vancouver, Canada) in order to obtain *bona fide* single-cell suspension. 1×10^3^ cells were plated in U-bottom 96-well plates and treated or not with 100, 200 and 400 µM Suramin. Cell growth was assayed at the indicated time using the CellTiter Assay System (Promega).

### Protein extraction and Western blot

The single-cell suspension was obtained by mechanical disaggregation of the spheres and total proteins were extracted 48 hours later, as already described (38). After separation by SDS– polyacrylamide gel electrophoresis, proteins were blotted on nitrocellulose membranes (GE Healthcare Europe Gmb) and hybridized with the following antibodies: anti-HMGA1 (70), anti-HMGA2 (Genetex) or affinity-purified anti-HMGA2 (71), anti-*α*-actin (SantaCruz Biotechnologies).

### RNA extraction and qRT-PCR analyses

The single-cell suspension was obtained by mechanical disaggregation of the spheres and RNA was extracted 48 hours later, by using Direct-zol RNA MiniPrep (Zymo Research). One μg of RNA was retrotranscribed, by using QuantiTect*®* Reverse Transcription Kit (Qiagen) and qRT-PCR was performed as described (38). Primers used in the qRT-PCR experiments are listed in Table S3. The 2^−ΔΔCt^ formula was used to calculate the differential gene expression (72).

### Wound Healing Assay

B16F10 melanoma cells were seeded at 10^4^ cells/cm^2^ in 6-well plates and treated with 100 µM of Suramin for 72h; then six scratches were made in confluent cultures using a 200 µL micropipette tip. Pictures were taken at 0 and 24h post-wounding at magnification of 40-fold. The invaded area was calculated using ImageJ software and the percentage invaded area was determined by calculating the invaded area at 24h to that at time 0h. Five fields of view per treatment were examined.

### Immunofluorescence

B16F10 melanoma cells were seeded at 10^4^ cells/cm^2^ in 24-well plates containing coverslips and treated with 100 µM of Suramin for 72h. Cells were then washed with PBS and fixed for 10 min with 4%PFA, permeabilized with 0.25% Triton X-100 and blocked for 1h with 10% goat serum. Subsequently, cells were incubated for 1h with Rabbit polyclonal IgG E-cadherin antibody (Santa Cruz, sc-7870) followed by incubation of Alexa 488 Goat anti-Rabbit secondary antibody for 1h. The staining was visualized using a Leica DMRB fluorescent microscope at 40x magnification. Pixel intensity was calculated using ImageJ software and the E-cadherin expression level was determined by calculating the pixel intensity of E-cadherin immunofluorescence staining over the total number of cells. Five random fields of view per treatment were examined.

### Molecular Dynamics Simulation

Since HMGA2 is shown to be an intrinsically disordered protein, we first performed molecular dynamics (MD) simulations to generate multiple disordered conformations of the suramin-interacting segment of HMGA2 chain (–GEKRPRGRPRKW–). Using the Charmm-Gui web interface(73), the peptide was solvated in a cubic water box with TIP3 water and the system was neutralized by adding five Cl^−^ ions. The final system contained ∼16,000 atoms. NAMD 2.12 (74) was used to perform all-atom molecular dynamics with CHARMM36 force field (75). The particle mesh Ewald (PME) method (76) was used for calculating the long-range ionic interactions. The system was minimized for 10,000 steps, followed by a 100 ps equilibration at 300K with 1 fs time step. A 100-ns production simulation with 2-fs time step was performed at a constant pressure of 1 atm. and T=300K. The Nose-Hoover Langevin-piston method (77) was used for pressure coupling, with a piston period of 50 fs and a decay of 25 fs, and the Langevin temperature coupling with a friction coefficient of 1 ps^-1^ was used for maintaining the temperature. From the 100-ns simulation trajectory, 1000 protein pdb frames were extracted using Visual Molecular Dynamics (VMD)(78).

### In silico docking studies

Suramin was docked to 1,000 MD-generated conformations using AutoDock Vina 1.1.2. (79). The protein pdb files and the suramin compound structure were first converted to pdbqt format for docking. Using custom scripts, Suramin was screened against the protein conformations and the resulting scores of the complexes were sorted and ranked according to their binding affinities.

## Acknowledgments

This work was supported by grants 1R15GM109254-01A1 and 1R21AI125973-01A1 from the National Institutes of Health (to F.L.), and by the Florida Translational Research Program, a contract administered by the Florida Department of Health (COHK8), and for which Layton Smith is the Principal Investigator.

## Author contribution

F.L., S.B., S.V., and L. Smith designed research; L.S., N.B. S.B., J.F., A.G., F.D., M.R., F.F. and P.C. performed research; F.L., S.B., A.F., L.K., F.F.-L., P.C., S.V., and L. Smith analyzed data; F.L. wrote the paper.

## Additional Information

Supplementary information accompanies this paper at *Nucleic Acids Research*.

