## Supplemental materials for "Suramin potently inhibits binding of the mammalian high mobility group protein AT-hook 2 to DNA"

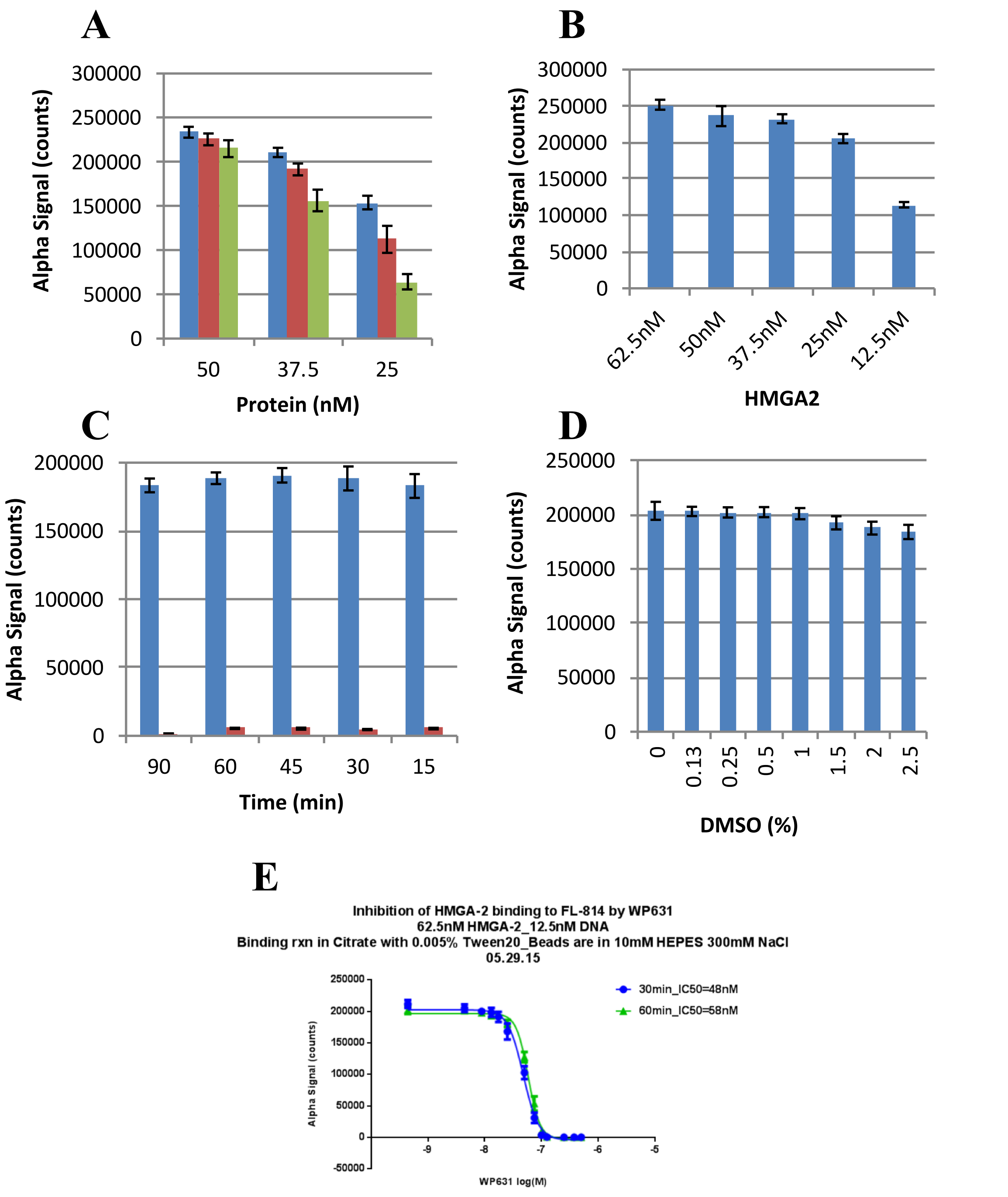


**Figure S1.** DNA-binding studies to determine the optimal conditions for HMGA2-FL814 interactions in 1×assay buffer (30 mM Citrate buffer, pH 7.0, 300 mM NaCl, 0.005% Tween 20) for the AlphaScreenprimary assay. (A) Varying HMGA2 and FL814 concentrations. Blue, red, and green columns represent 12.5, 6.25, and 3.125 nM of FL814. (B) Different concentrations of HMGA2 titrating into 12.5 nM FL814. (C) Time course of HMGA2-FL814 interactions. Blue and red columns represent the reactions in the presence and absence of HMGA2, respectively. (D) DMSO tolerant assays. (E) WP631 strongly inhibits HMGA2-FL814 interactions.

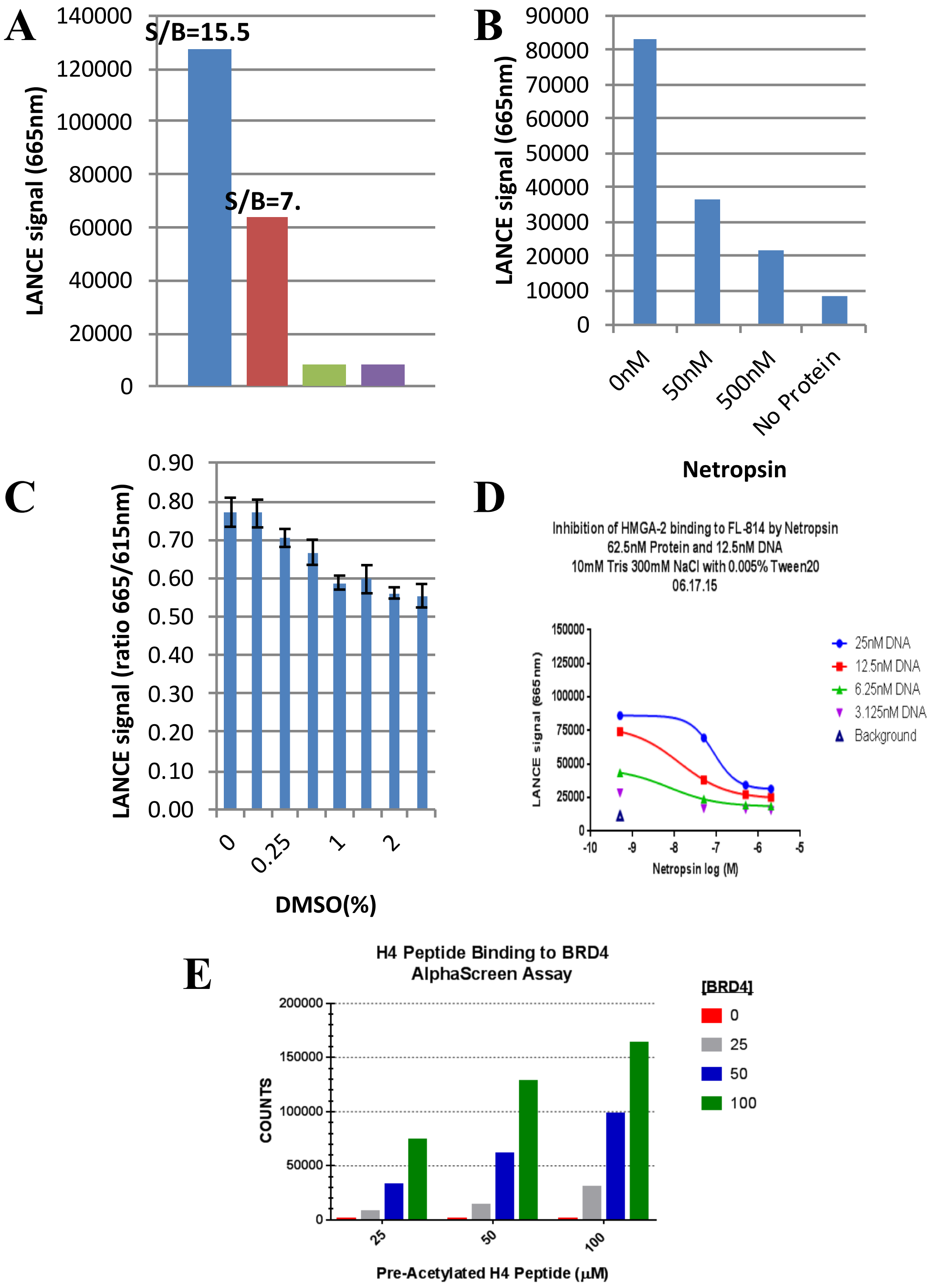


**Figure S2.** (A-D) DNA-binding studies to determine the optimal conditions for HMGA2-FL814 interactions for the LANCE time-resolved fluorescence energy transfer (TR-FRET) assay. (A) Varying HMGA2 and FL814 concentrations. Blue column: 62.5 nM HMGA2 and 12.5 nM FL814. Red column: 31.25 nM HMGA2 and 6.25 nM FL814. Green colun: 12.5 nM FL814 only. Purple column: 6.25 nM FL814 only. (C) DMSO tolerant assays. (B) and (D) Netropsin strongly inhibits HMGA2-FL814 interactions. (E) The AlphaScreen assay for H4 peptide binding to BRD4.

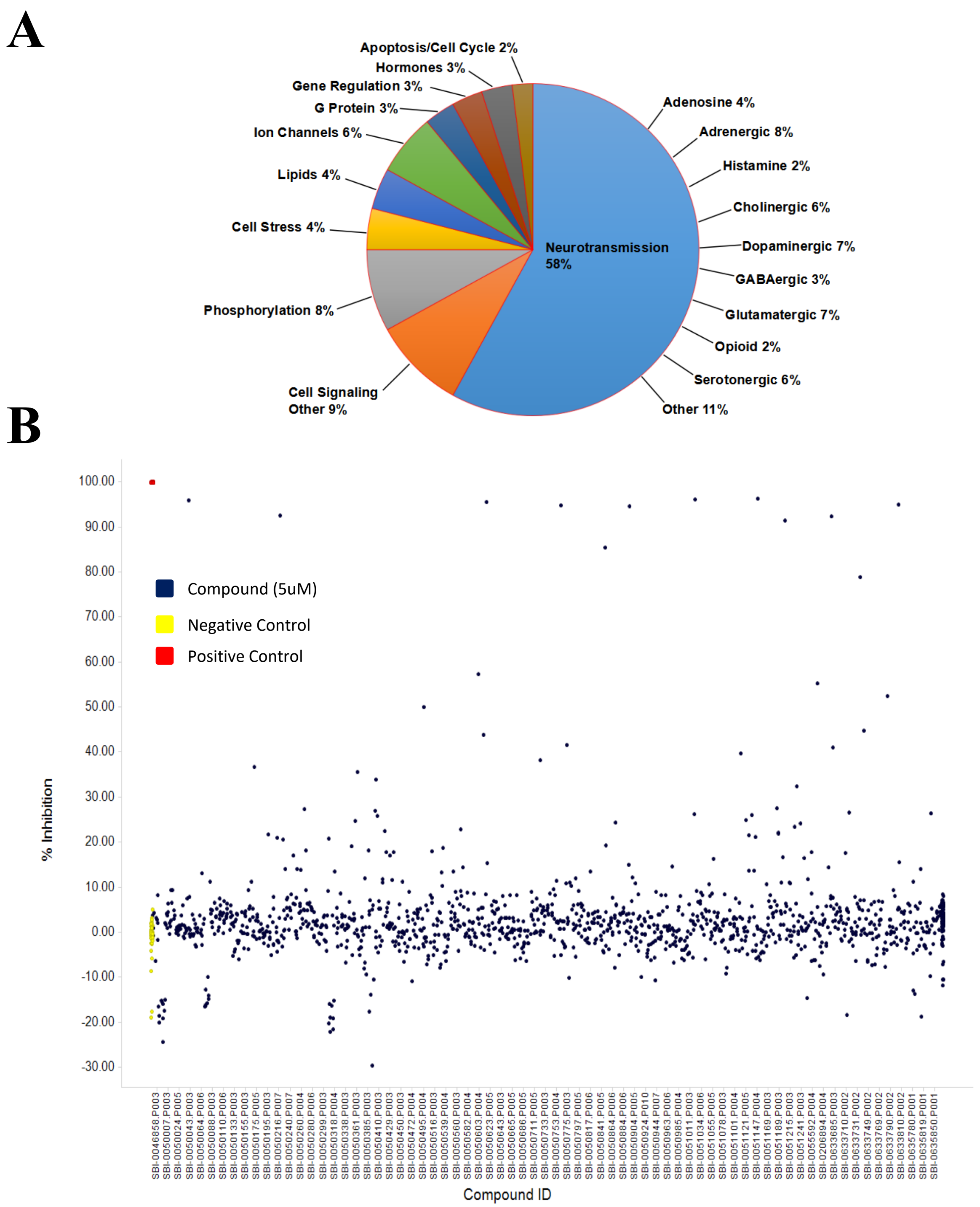


**Figure S3.** (A) The biologically annotated collection of LOPAC1280 compound library that contains 1280 pharmacologically active compounds. (B) HMGA2-DNA pilot screens using the Sigma LOPAC1280 compound library. Netropsin was used as positive controls.

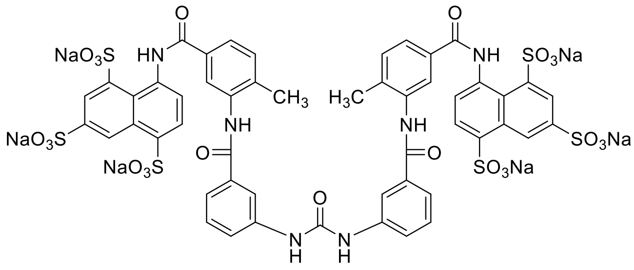

**Suramin**

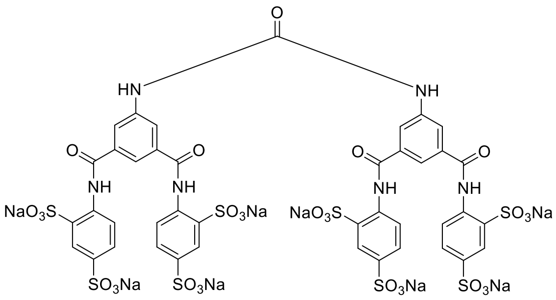

**NF449**

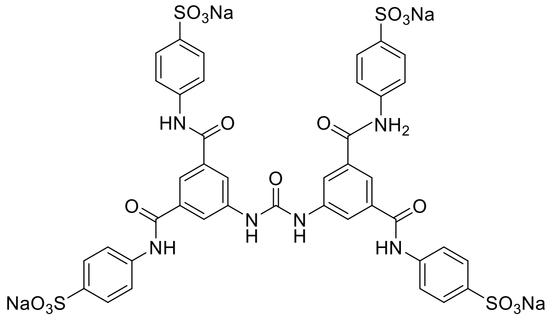

**NF110**

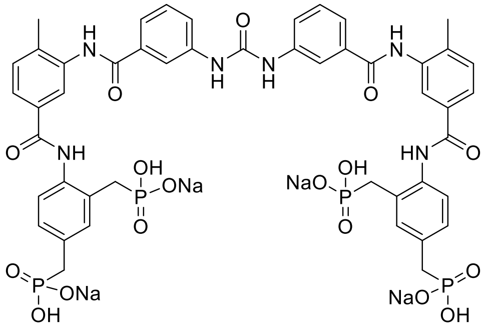

**NF546**

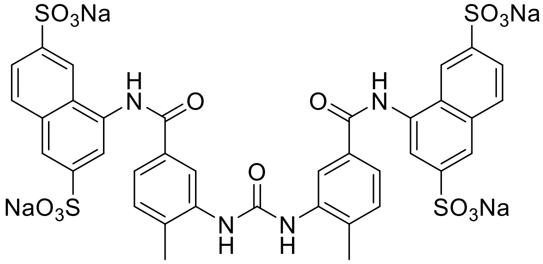

**NF340**

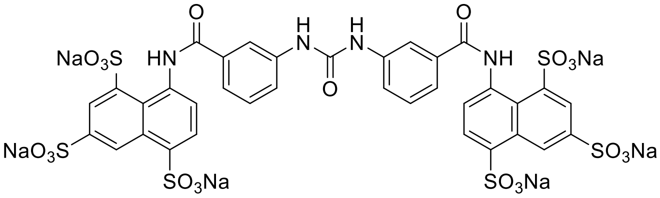

**NF023**

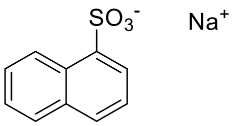

**N0015**

**Figure S4.** Chemical structures of suramin and analogues.

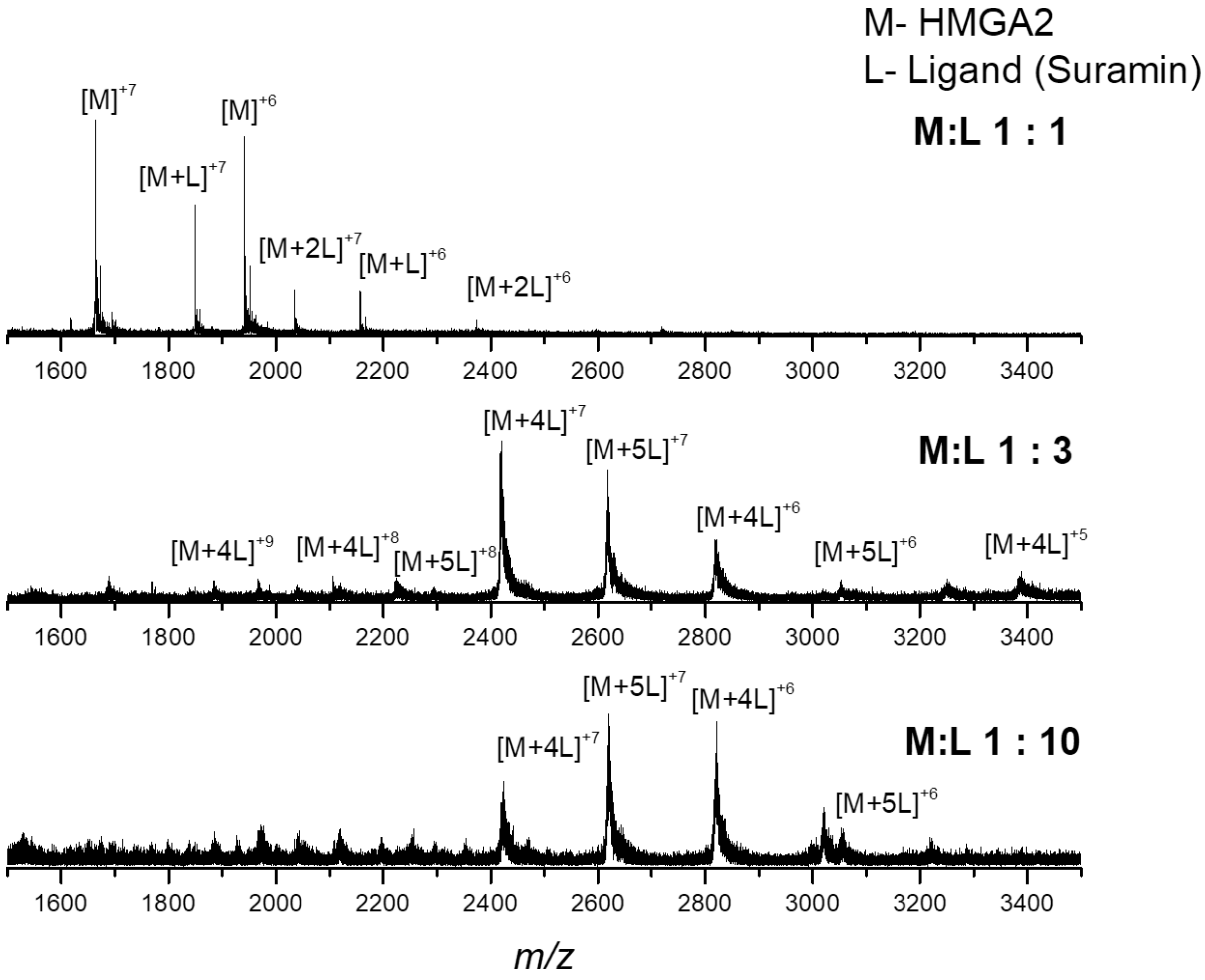


**Figure S5.** Typical native mass spectrometry spectra of a mixture of HMGA2:Ligand (Suramin) at 1:1, 1:3 and 1:10 ratio. Notice the increase in the numbers of ligands up to [M+4L] and [M+5L] per HMGA2 molecule observed in the MS from 1:1 to 1:10 ratios.

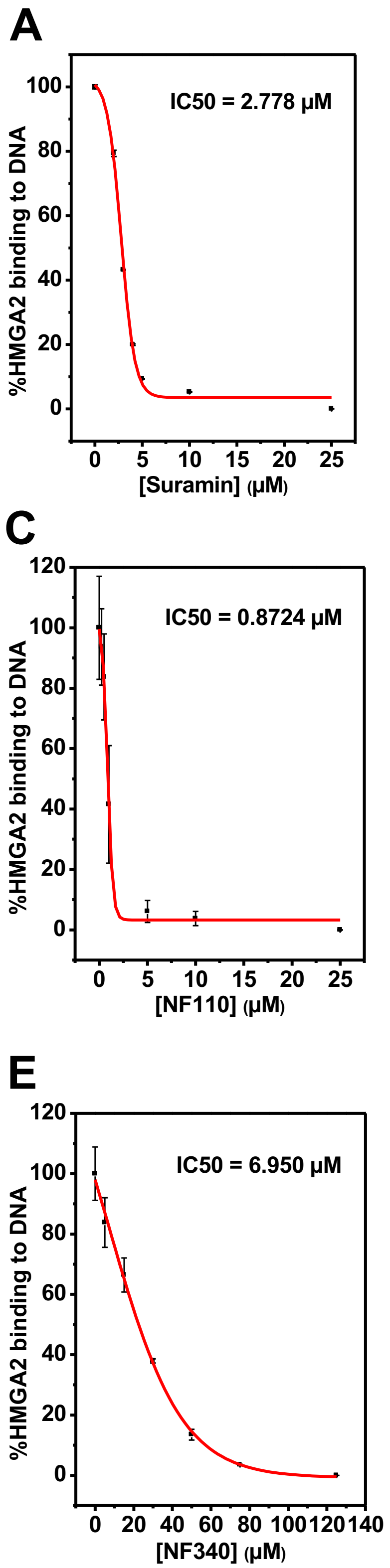


**Figure S6.** Inhibition of HMGA2 binding to FL814 by suramin (A) and analogues (B-F). The apparent inhibitory IC50 was determined by using PDI-ELISA.

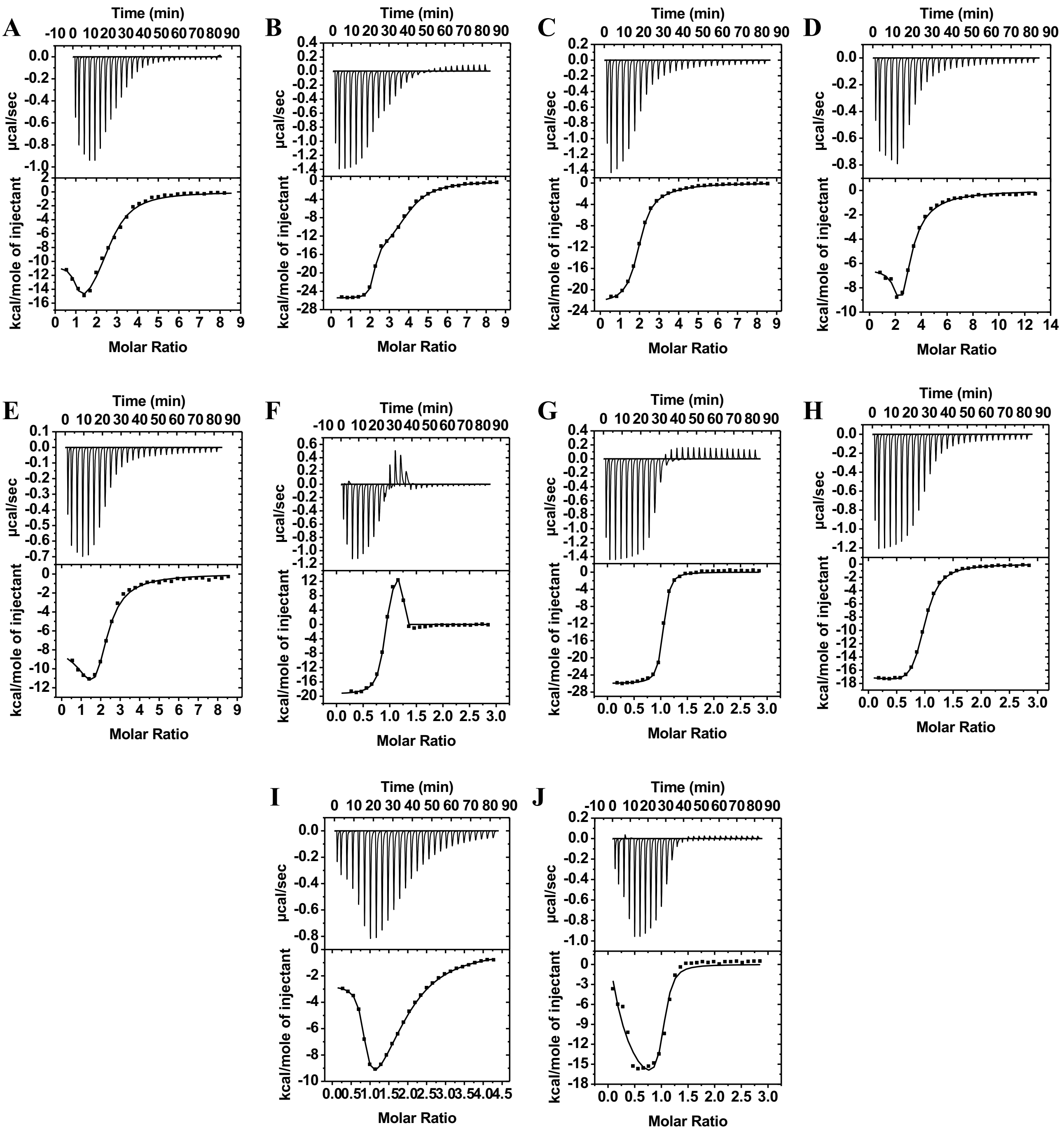


**Figure S7.** Sample raw data from isothermal titration calorimetry (ITC) experiments for the titration of suramin analogues to HMGA2 (A-E) and ATHP3 (F-J). ITC experiments were performed according to conditions as described in Materials and Methods. The ITC data were fit using the software supplied by the manufacturer to yield thermodynamic parameters. (A) and (F) NF449; (B) and (G) NF110; (C) and (H) NF546; (D) and (I) NF 340; (E) and (J) NF023.

.

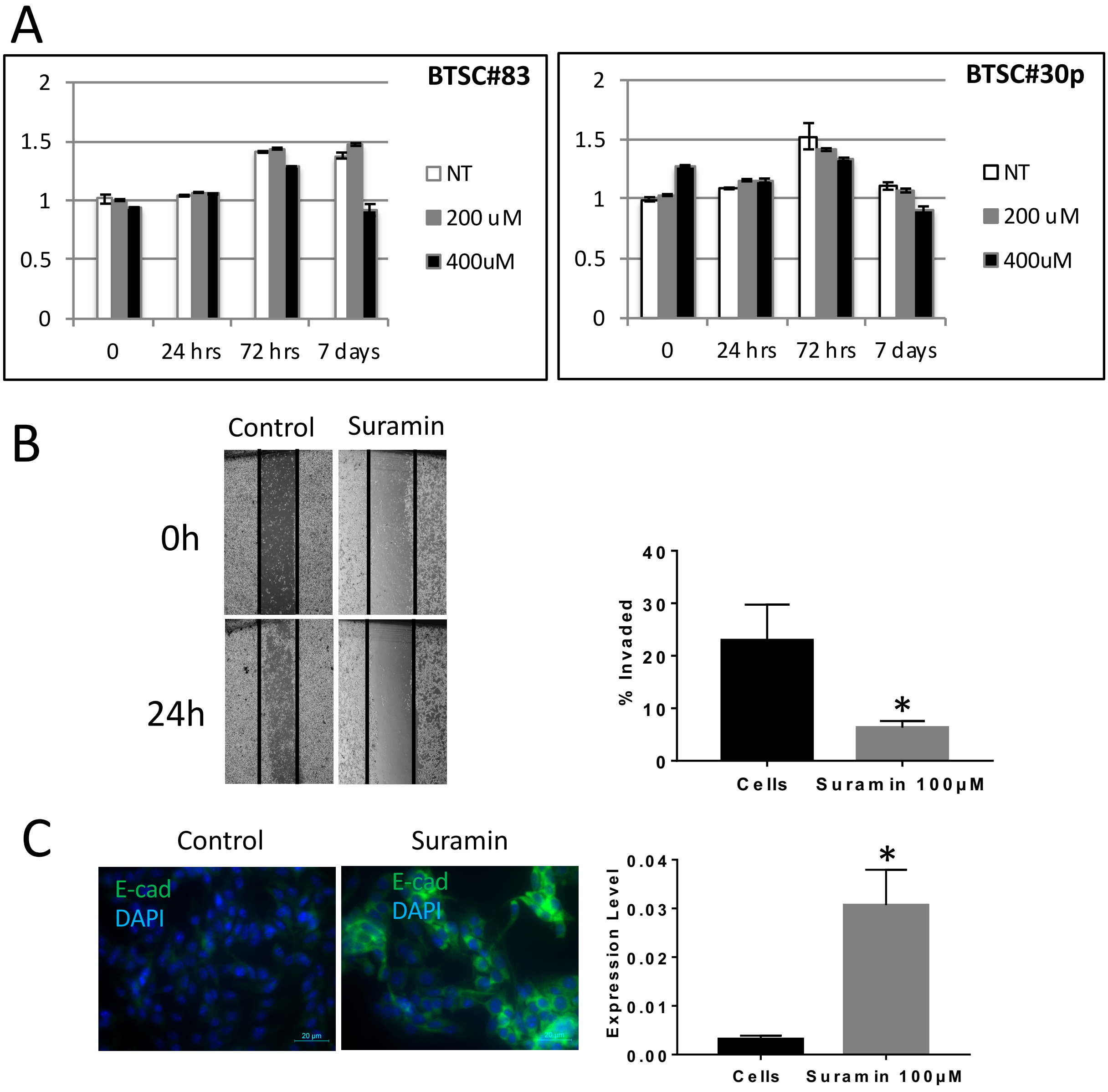


**Figure S8.** (A) Growth curves of BTSC#83 (left panel) and BTSC#30p (right panel) grown in U-bottom 96-well plates in presence of 200 and 400 µM of suramin. Means and standard deviations are calculated from three independent experiments. (B) Suramin decreases the migratory behavior. Left panel: representative images photographed at 0 and 24h post-wounding shown at magnification of 40x. Right panel: quantitative analysis of migratory behavior of B16F10 melanoma cells treated with Suramin at 100µM for 72h. Percentage Invaded area was determined by calculating the invaded area at 24h to that at time 0h. Five fields of view per treatment were examined. Data were represented as mean ±SEM ($p<0.05$). (C) Increases E-cadherin expression of B16F10 melanoma cells. Left panel: representative images of E-cadherin immunofluorescence staining shown at magnification of 40x. Right panel: the expression level of E-cadherin was determined by calculating the pixel intensity of E-cadherin immunofluorescence staining over the total number of cells. Five random fields of view per treatment were examined. Data were represented as mean ±SEM ($p<0.05$).

**Table S1.** Parameters for the primary HTS assay of HMGA2-DNA interactions using LOPAC library.

| | |
|---|---|
| # of compounds | 1280 |
| Tested concentration | 5 μM |
| Z' value | 0.83 |
| RZ' value | 0.9 |
| [a]S/B | 438 |
| # of compounds with inhibition>50% | 17 |
| Hit rate | 1.25% |

[a]S/B represents signal versus background ratio.

**Table S2. Inhibitors identified by the HTS assays**

| Compounds | IC50 (μM) | | | |
|---|---|---|---|---|
| | Alpha Screen | Lance Screen | BRD4 | Cytotoxicity |
| WP631 | 0.044 | 0.17 | 2.05 | 0.17 |
| Cisplatin | > 100 | 9.00 | > 100 | > 100 |
| cDPCP[a] | > 100 | 15.41 | > 100 | > 100 |
| Ro 90-7501[a] | 1.32 | 11.58 | 16.90 | > 100 |
| ATA[a] | 0.32 | 0.21 | 5.00 | > 100 |
| Mitoxantrone | 0.18 | 1.77 | 0.99 | 1.65 |
| Suramin | 4.06 | 2.58 | > 100 | > 100 |
| Carboplatin | > 100 | > 100 | > 100 | > 100 |

[a]cDPCP, cis-Diammine(pyridine)chloroplatinum(II) chloride; Ro 90-7501, 2′-(4-Aminophenyl)-[2,5′-bi-1H-benzimidazol]-5-amine; ATA, aurintricarboxylic acid.

**Table S3. DNA oligonucelotides used for primers of the RT-PCR experiments**

| Oligo names | Sequence (5-3’) | Gene |
|---|---|---|
| HMGA1_F[a] | CAACTCCAGGAAGGAAACCA | HMGA1 |
| HMGA1_R[a] | AGGACTCCTGCGAGTGC | HMGA1 |
| HMGA2_F | TCCCTCTAAAGCAGCTCAAAA | HMGA2 |
| HMGA2_R | ACTTGTTGTGGCCATTTCCT | HMGA2 |
| SNAIL_F | AGTGGTTCTTCTGCGCTACT | SNAIL |
| SNAIL_R | GGGCTGCTGGAAGGTAAACT | SNAIL |
| TWIST_F | TCTCGGTCTGGAGGATGGAG | TWIST |
| TWIST_R | GTTATCCAGCTCCAGAGTCT | TWIST |
| NUMB_F | ACAGAGAGGCACGCTATGCT | NUMB |
| NUMB_R | TAGCTCTGCTGGGCAGTTG | NUMB |
| ID2_F | TGTCAAATGACAGCAAAGCAC | ID2 |
| ID2_R | GTTGTTGTTGTGCAAAGAATAAAAG | ID2 |

[a]F and R represent forward and reverse orientations, respectively.
